# Transcriptomics data availability and reusability in the transition from microarray to next-generation sequencing

**DOI:** 10.1101/2020.12.31.425022

**Authors:** Gabriella Rustici, Eleanor Williams, Mitra Barzine, Alvis Brazma, Roger Bumgarner, Marco Chierici, Cesare Furlanello, Liliana Greger, Giuseppe Jurman, Michael Miller, B.F. Francis Ouellette, John Quackenbush, Michael Reich, Christian J. Stoeckert, Ronald C. Taylor, Stephen Chervitz Trutane, Jennifer Weller, Brian Wilhelm, Neil Winegarden

## Abstract

Over the last two decades, molecular biology has been changed by the introduction of high-throughput technologies. Data sharing requirements have prompted the establishment of persistent data archives. A standardized approach for recording and managing these data was first proposed in the Minimal Information About a Microarray Experiment (MIAME) guidelines. The Minimal Information about a high throughput nucleotide Sequencing Experiment (MINSEQE) proposal was introduced in 2008 as a logical extension of the guidelines to next-generation sequencing (NGS) technologies used for transcriptome analysis.

We present a historical snapshot of the data-sharing situation focusing on transcriptomics data from both microarray and RNA-sequencing experiments published between 2009 and 2013, a period during which RNA-seq studies became increasingly popular for transcriptome analysis. We assess how much data from RNA-seq based experiments is actually available in persistent data archives, compared to data derived from microarray based experiments, and evaluate how these types of data differ. Based on this analysis, we provide recommendations to improve RNA-seq data availability, reusability, and reproducibility.

## INTRODUCTION

Microarray technologies significantly changed biology and medicine, by providing one of the first platforms for assessing gene expression levels on a genome-wide scale simultaneously, and the so-called SNP-chips (single-nucleotide polymorphism microarrays) added the dimension of common variant detection within the transcripts. Next-generation sequencing (NGS) technologies provide additional dimensions of information that have proven to be of great clinical value, allowing detection of isoforms [1, 2], allelic balance [3], editing [4–7] and rare variants, but the NGS platforms present very similar data volume, storage, and meta-data reporting challenges to microarrays.

Sample preparation methods for high-throughput technologies are complex and greatly influence both transcript profiles and interpretation of the resulting measurements. Experimental design must be sophisticated and matched with appropriate pre-processing and statistical data analysis before interpretations are accepted as valid: the need to report analysis workflows in the same detail as sample preparation protocols first became apparent for microarray experiments. The results of highly parallel experiments are particularly valuable when small studies are combined into larger meta-analyses, but the factors conditioning individual studies must be explicitly detailed to enable the design of valid meta-analyses. In order to maximize the evaluation and reuse of primary research data, these data and the derived information must be shared through persistent public archives. The very large size of the raw and processed data sets requires a considerable investment in research infrastructure as well; both economy and a desire to promote data integration encouraged the creation and sharing of ontologies and data representation standards. Thus functional genomics scientists were early proponents of the need for workflows, ontologies and sharing raw and derived data.

Guidelines for insuring that a microarray experiment and its results will remain interpretable were published in 2001 as the Minimum Information About a Microarray Experiment (MIAME) [1]. With the shift to the newer NGS technology it was natural that scientists who had used microarrays would expect a similar reporting discipline for their NGS experiments. In 2012, the Functional Genomics Data (FGED) Society (http://www.fged.org/) published a revised version of the Minimum Information about a high-throughput nucleotide SEQuencing Experiment (MINSEQE) guidelines (http://www.fged.org/projects/minseqe/). MINSEQE guidelines are similar to MIAME guidelines, consisting of the following main elements:

1. Experimental design and sample-data relationships: this includes a summary of the experiment and its goals, researcher contact information, identifiers for associated publications, and specifying the relationships between samples and data files.
2. A full description of the biological system and treatments used, including how samples derive from the organism and its tissues, and the experimental variables being studied (e.g. “compound” and “dose” in dose-response experiments).
3. Essential laboratory experimental and data processing protocols, including sample isolation, RNA purification and processing prior to sequencing or hybridization, the preparation strategy for making a sequencing library or labelling a transcript pool, any internal labeling (such as bar codes) and amplification methods; data pre-processing (such as alignment for sequencing or filtering for significant signal levels on microarrays), data processing and data analysis protocols.
4. The ‘raw’ data (CEL or GPR files for microarrays and sequence read data for NGS) for each assay: primary derived data (spot-intensity data or sequence reads and base-level quality scores) for each assay. Recommendations concerning standard reporting formats may be suggested, such as the FASTQ format for sequence data and the scale used for quality scores.
5. The ‘final’ processed (or summary) data for the set of assays in the study: the data on which the conclusions in the related publications are based, and descriptions of the data formats.
6. For microarrays a separate category specifies detailed microarray design features and architecture, including the probe sequences and their placement.

A number of public data repositories have adopted and implemented the MIAME guidelines, and provide full access to microarray and NGS data; the largest and most highly referenced databases include ArrayExpress at the European Bioinformatics Institute (EBI, http://www.ebi.ac.uk/arrayexpress/) [2], the Gene Expression Omnibus (GEO) at the National Center for Biotechnology Information (NCBI, http://www.ncbi.nlm.nih.gov/geo/) [3], and the Omics Archive at the DNA Data Bank of Japan (DDBJ, http://trace.ddbj.nig.ac.jp/) [4]. Other repositories include such meta-data but limit access due to ethical and legal restrictions on human research subjects, such as the database of Genotypes and Phenotypes (dbGaP) at NCBI [5] (http://www.ncbi.nlm.nih.gov/gap) and the European Genome-phenome Archive (EGA) at the EBI (https://www.ebi.ac.uk/ega/)[6]. However, the public aggregate information and non-identifiable processed data from studies with controlled access data in dbGAP and EGA can be browsed on public websites which provide information about the aim, the experiments and the data used in the registered studies but data access to individual studies is subjected to approval from a Data Access Committee. Most major scientific journals now require submission of microarray data to a MIAME-compliant and persistent archive of the data as part of the publication process.

In order to cope with the explosive growth of NGS data and the associated storage needs, an international public archival resource, the Sequence Read Archive (SRA) [7] was established in 2009, as a collaboration between NCBI, EBI and DDBJ. SRA is used for archiving and sharing of all kinds of raw sequencing data, including functional genomics studies. Data may be deposited in the archives prior to publication to comply with a journal’s data transparency requirements, or as a stand-alone data release in the absence of a peer-reviewed publication. An alternative route to direct SRA submission is to go through ArrayExpress, GEO or the Omics Archive, that act as ‘brokers’, depositing the associated raw data files in the SRA while retaining some meta-data and processed data, such as gene expression levels. These repositories have submission systems in place to enable capture of required meta-data attributes.

In order to assess the level of compliance and the usability of data in these repositories, we used a combination of experts in the field and software scripts to characterize a subset of available data. In carrying out our studies, we mined the Europe PubMed Central (Europe PMC) [8] for articles reporting the results of gene expression experiments using either microarray or RNA-seq data, or both. Only open access data was included in the analysis, i.e., the restricted-access data from EGA and dbGAP was excluded. Articles were linked to data deposited in three primary repositories: ArrayExpress, GEO and SRA. We focused our analyses on microarray and RNA-Seq data as most directly comparable, but believe similar trends will be seen for ChIP-Seq and ChIP-chip data. We examined MIAME and MINSEQE compliance for the selected data sets to determine the granularity of the reported experimental meta-data in each, and to determine if there are systematic differences between them. Although full reanalysis of published results is not possible in such a survey, we attempted to determine if the repositories capture sufficient information to permit effective reuse of the data.

## RESULTS

The results of our analysis of submission rates (see Table 1) show that ~65% of the publications we reviewed in the period 2009 - 2013 containing new RNA-seq data have associated data submitted to ArrayExpress, GEO or SRA. By comparison, the data submission rate for microarray data for one of those years (2012) was ~ 49%, a value similar to that reported by Piwowar in 2011 (45%) [9].

**Table 1.** Percentage of RNA-seq publications from 2009 to 2013 containing new gene expression data from RNA-seq experiments in which the data has been submitted to ArrayExpress, GEO or SRA. To permit comparison to microarray publications (a much larger data set), percentages were calculated for microarray gene expression data in 600 randomly selected publications from 2012, comparable to the 537 RNA-seq publications available for that year.

| Type of data |  | 2009 | 2010 | 2011 | 2012 | 2013 |
| --- | --- | --- | --- | --- | --- | --- |
| RNA-seq | Number of publications with new data | 10 | 76 | 173 | 332 | 523 |
| RNA-seq | % publications with data submitted to databases | 70 | 67 | 68 | 61 | 65 |
| Microarray | Number of publications with new data |  |  |  | 347 |  |
| Microarray | % publications with data submitted to databases |  |  |  | 49 |  |

Of experiments published in 2012 and 2013 based on RNA-seq datasets for which no database accession could be identified, only 13% and 15% respectively were studies whose samples came from human individuals (see Table 2). This suggests that there are reasons other than ethical issues as to why these data were not submitted to public archives. The articles associated with such datasets were published in a wide range of journals, and no bias was observed between “non-submitted” data and a particular scientific journal or editorial group (data not shown).

**Table 2.** The type of samples used in RNA-seq publications in 2012 and 2013 that were not submitted to ArrayExpress, GEO or SRA.

| Year | % publications with human individual samples | % publications with human cell line samples | % publications with non-human samples | Number of journals |
| --- | --- | --- | --- | --- |
| 2012<br>(126 total publications) | 13 | 5 | 82 | 63 |
| 2013<br>(184 total publications) | 15 | 12 | 73 | 106 |

### MINSEQE compliance

The quality of the meta-data and the level of biological replication are of crucial importance to the suitability of data for re-use [10]. We compared the presence of experimental meta-data for 100 RNA-seq datasets each across ArrayExpress, GEO and SRA, using compliance with the MINSEQE guidelines as our metric. The results of this analysis are summarized in Table 3. It is evident that the three databases differ in the way they request, record and display the meta-data associated with RNA-seq experiments. In general, ArrayExpress and GEO, which have implemented the MIAME guidelines for microarrays in the past, more effectively capture RNA-Seq experimental details and protocols. For example, experimental variables, which are the core of any statistical analysis of the data, and with sample type and replication will determine re-use of the data, are present in 98% of ArrayExpress and GEO entries but only 74% of SRA entries. An even larger difference was found for the presence of processed data (e.g. BAM files or FPKM tables) in our selected experiments, available in 91% of GEO entries, 24% of ArrayExpress entries and 0% of SRA entries.

**Table 3.** Percentage of RNA-seq studies providing meta-data, protocols, raw and processed data files from 100 randomly selected datasets in ArrayExpress, GEO and SRA.

|  | ArrayExpress | GEO | SRA |
| --- | --- | --- | --- |
| Aim of experiment | 98 % | 100 % | 91 % |
| Type of experiment | 100 % | 100 % | 100 % |
| Contact details | 100 % | 100 % | 0% |
| Sample information (species) | 100 % | 100 % | 100 % |
| Experimental variable (factor information) | 99 % | 97 % | 74 % |
| Biological and technical replicates | 93% | 80 % | 86 % |
| Treatment protocol if applicable | 100 % | 100 % | 34 % |
| Library construction protocol | 85 % | 87 % | 36 % |
| Raw sequencing reads | 99 % | 98 % | 100 % |
| Processed data | 24 % | 91 % | 0 % |
| Data processing protocol with information about software and genome build for read mapping (where processed data present) | 67 % | 81 % | 0 % |
| % studies that meet MINSEQE standard | 14% | 62% | 0% |

While completeness of the meta-data was our primary focus, during the analysis we often observed that even when the meta-data for a given study was available, it is was not reported in a consistent manner. For example, sometimes the experimental factor was present as part of the sample title, in other cases it was part of the sample description, or it might be in the sample attributes. In some cases detailed meta-data is only to be found embedded in XML files, and the researcher must know how to retrieve and inspect them when filtering for relevant experiments. The authors are bioinformaticians familiar with these types of data and these repositories, and even so, a user familiar with the style of one interface could be misled by the way the information is presented through a different interface, or how it is stored in supplementary files, thereby missing relevant information. This became clear for the part of the study in which two assessors looked at the same files. There was agreement for most essential information such as the type of experiment (100% agreement), the species targeted (100%) or the processed data availability (100%), the level of agreement began to decline when asked about information explaining experimental variables (82%), experimental treatments (69%), biological replicates (73%), library preparation methods (78.3%) or data processing protocols (78%). This decline suggests that the information is both hard to find and that there is disagreement in what level of detail the information protocols should include. Agreement was higher when retrieving information from ArrayExpress (92%) and GEO (90%) compared to SRA (80%). This discrepancy might be in part due to the fact that two different user interfaces are available for the SRA, which display information differently, leading to possible disagreement.

### Data reuse

In the course of our analysis we encountered a number of publications reusing existing RNA-seq data, either as part of their interpretation of new experimental data, or to compare with findings based on analysis of microarray data [11]. Throughout 2013, the reused datasets had an average MINSEQE score of 3.7 (where a MINSEQE score of 5 indicates full compliance with the guidelines) where the lack of a higher score came primarily from inadequate or missing protocols (present in 50%) or missing processed data (present in only 41.25%). This suggests that the availability of raw data (95%) and experimental variables (87.5%) are considered acceptable indices for reusability. This trend is also seen with the Expression Atlas at EBI, which reanalyzes RNA-seq data, from ArrayExpress and other sources, to present information about the properties of genes or samples in both baseline and differential expression estimates [12]. The availability of raw data and clearly defined experimental variables were again the two most important criteria for re-use of datasets, but it can be seen that they are also filtered on protocol information (to ensure appropriate re-analysis methods are used) and for a minimum number of 3 biological replicates (for statistical confidence estimates). The outcome is that less than 10% of RNA-seq submissions to ArrayExpress are suitable for inclusion in the Expression Atlas.

The property of biological replicates has been heavily emphasized in reviews of the microarray gene expression literature. We further investigated this aspect of our data, comparing the number of biological replicates between ~100 RNA-seq and microarray studies matched for equivalent experimental conditions (Figure 1). On average, RNA-seq studies have 2.8 replicates, compared to 3.6 in microarray studies, likely due to the higher cost of the technology at that time, and larger sample mass needs. On the no-replicates end of the spectrum, 27% of RNA-seq studies fall in this group versus 9% of microarray experiments. The inevitable outcome, since a minimum of 3 replicates is required for calculating the standard deviation, is that about one quarter of RNA-seq studies from this period are extremely limited in the type of statistical analysis that can be performed to determine differential expression with adequate confidence levels.

**Figure 1.**
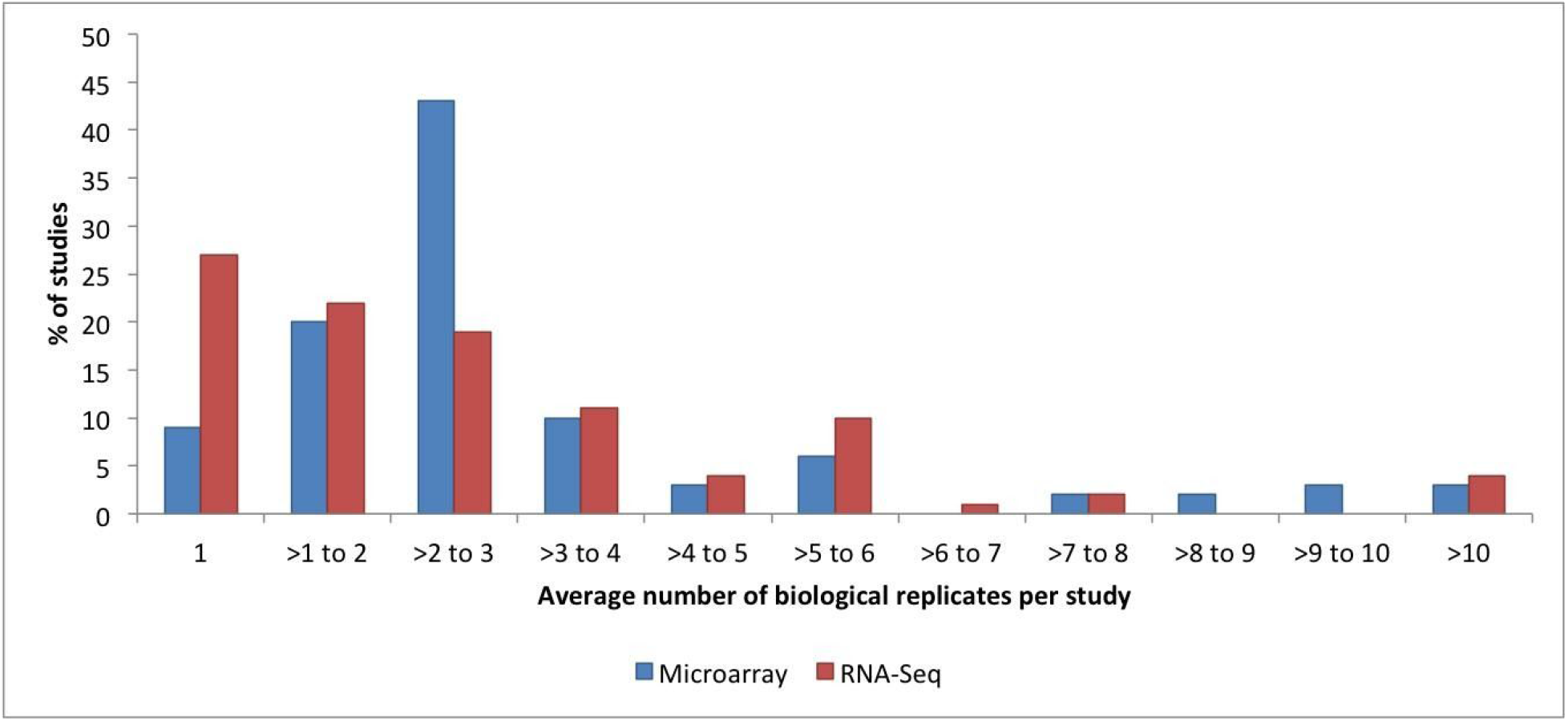
The average number of biological replicates per microarray and RNA-seq study in ArrayExpress and their frequencies.

## DISCUSSION

We found it encouraging that the overall availability of the data from NGS based functional genomics experiments is higher (65%) than that for microarray experiments (49%). This may seem surprising given that the MIAME initiative is 10 years older, but it may follow from the even longer history for depositing DNA sequences to international consortium databases, which dates back to 1982 for GenBank (http://www.ncbi.nlm.nih.gov/genbank/statistics), and the policy of the Human Genome Project requiring the release of sequence data within 24 hours of its production (known as the Fort Lauderdale and Toronto agreements).

The MIAME standard went considerably beyond recommendations for simple deposition of data. It included the need for standard data formats and complete, interpretable meta-data sufficiently detailed to allow reproduction of the quite complicated experiments. Assessing data reproducibility and general reusability is challenging. In 2009 Ioannidis and colleagues [17] assessed the reproducibility of results from 18 microarray-based studies for which data had been deposited in GEO or ArrayExpress between 2004 and 2005. The results were quite worrying – only a minority of the results could be reproduced; assuming that the published results were correct, it was concluded that the lack of reproducibility was largely due to the unavailability of necessary meta-data. We did not undertake to check the reproducibility of the results reported for the NGS functional genomics studies we examined, instead taking the availability of essential meta-data as a proxy for data reusability.

A conclusion of this study is that for those data sets directly submitted to the SRA, meta-data important to data interpretation were not available. However, HTS data sets brokered by GEO and ArrayExpress have been annotated to about the same level of detail as the available microarray data. GEO has considerably more processed data than ArrayExpress, while ArrayExpress has more detailed sample annotation: we think that both are needed for reusability. Clearly, an experiment makes little sense if the essential sample properties and experimental variables are not provided. On the other hand, given the complexity of RNA-seq analytical workflows, it is beneficial to include both raw and processed data files associated with an RNA-seq study. Processing methods for RNA-seq data are still evolving, so providing output for defined checkpoints in an analysis pipeline allows a common starting point for testing alternate or new analysis methods without having to revert to processing raw data, which requires expertise and computing infrastructure available to only a few researchers. A BAM file containing the mapped reads or a table of counts containing the expression levels estimates for genes or transcripts provides an excellent starting point for additional or meta analyses.

Data deposition levels are at ~50% and 65% respectively for microarray and NGS publications. Initially we hypothesized one reason for the lack of data availability was due to use of human research subjects. However, our analysis showed that this not the case for the studies we selected, which had quite low numbers of studies including human subjects. Where such data are generated by large consortia, such as the International Cancer Genome Consortium (ICGC), there is a strict data release policy to a protected database that we did not access, so our conclusions about annotation compliance cannot be extended to those archives.

With respect to the archives themselves, the SRA is specialized for the storage of large volumes of sequence data. Our research clearly showed that for studies that have been directly submitted to SRA, as opposed to brokered via GEO or ArrayExpress, meta-data important for interpreting the studies were not available. Since ~80% of RNA-Seq data in the SRA is not so brokered, a large majority of the experiments likely cannot be replicated and should not be reused.

We also wonder how it is possible that after more than 10 years of the MIAME initiative, 51% of publications relying on microarray data do not provide it in the permanent archives? This can be viewed as a significant failure of the community and scientific journals and their reviewers in particular. If reinterpretation and incorporation into meta-experiments is an important function for these data then the MIAME and MINSEQE initiatives remain unrealized and important goals. In addition, a high proportion (approximately 80%) of RNA-seq data in SRA have been submitted directly and did not come through ArrayExpress or GEO and is therefore likely to be under-annotated and less likely to be repeatable or re-usable.

It is also possible that some of the NGS experiment meta-data is unavailable because of how experiments are often performed: researchers often carry out the experimental design through sample acquisition and processing in their own labs, but then send the RNA (or cDNA) to a large sequencing center for subsequent parts of the workflow, and perhaps data goes to a separate genomics analysis group for early steps in the analysis workflow. Data submission responsibility usually lies with one of the latter groups. Genome sequencing centers often have well established data submission pipelines to SRA so their part of the annotation is quite complete, but they do not have methods for acquiring and incorporating the sample meta-data, nor subsequent analysis workflows. The solution to this should be in developing and using data management and annotation tools that can be shared with the originating labs, allowing them to record all the steps as the samples are processed, and similar tools for the Genomics Analysis groups, to capture the workflows and data at specified checkpoints.

We feel that the adoption of best practices for reporting details of analysis pipelines and ultimately the source code associated with a scientific publication is the only way to make an analysis fully reproducible [13]. Open source software tools are a potential key to solve this issue. Projects like Bioconductor [14], Galaxy [15] and GenePattern [16] allow reporting of the exact version of the software used so that discrepancies between versions can be tracked, give access to the source code being used and document all the analytical steps, ensuring transparency and reproducibility. Although we recognize that source code availability does not address whether the data was analyzed correctly or not, it will surely help to reduce uncertainties about a study’s conclusions and readily allow testing for reproducibility.

In the future, further shaping of the MINSEQE guidelines is likely to be necessary, as the technology and its biological applications evolve. Current specifications may become redundant or irrelevant while new information will need to be added. Examples of new specifications might be the requirement to include adapter sequences for small RNA study submissions and spike-in sequences for single cell RNA-seq study submissions.

## MATERIALS AND METHODS

### Availability of published data in persistent data archives

#### I. RNA-Seq subset

Europe PMC [8] was searched for journal articles or letters published between 2009 and 2013, containing the keyword “RNA-seq”. Europe PMC is a full-text article database that extends the functionality of the original PubMed Central repository; among other features it indexes database accession numbers (such as ENA and ArrayExpress) present in journal articles, using text-mining approaches to find literature-database relationships [17].

Publications were selected based on full text accessibility to the authors, either through open access policy or journals’ subscription (92-100% depending on the year). The articles were manually classified into those that had newly-generated RNA-seq data associated with them and those that did not, the latter group included review articles, publications about new analysis methods tested on publicly available datasets and research articles using data previously generated. Articles referencing new RNA-seq data were checked for database accession numbers using a template-based search. Where a database accession number was not included in the associated article we cross-referenced PubMed identifiers to the databases to identify those for which there were submitted data sets. Where no accession number could be found we postulated two possibilities: no submission or human subject data requiring deposition in a controlled access repository. To check these possible outcomes we classified samples in the categories ‘human, individual’, ‘human, cell-line’ and ‘non-human, individual’ in the subset of publications occurring in 2012-2013 for which new RNA-seq data was reported but no database accession identifier could be found in the publication.

#### II. Microarray subset

To compare the submission rate of gene expression microarray data to that of RNA-seq data, we again used Europe PMC and searched for journal publications containing the key words “microarray” and “gene” and “expression”. A very large number of articles resulted so the set was limited to those published in 2012, which still left us with 5128 publications, too many for manual inspection. Therefore 600 were randomly selected for further analysis, comparable to the 537 RNA-seq publications available for 2012. The set was classified into the group of 347 that produced new, accessible gene expression microarray data, while the remainder did not. Again, the articles were manually searched for database accession numbers and cross-referenced with PubMed identifiers from these databases.

### Data Reusability Factors: Number of biological replicates

Sample type and study factor are the primary reasons to select experiments for meta-studies, and these features tend to be clearly annotated. However, studies with few biological replicates are of limited use in meta-analyses (or other types of re-interpretation) given that most statistical methods require at least 3 biological replicates to allow calculation of the degree of variability (standard deviation) between samples or to set common baselines across the individual experiments. There has been a consistent message to researchers about the required level of replication for microarray experiments, but that message has been less emphatic for RNA-seq experiments. As part of this survey we examined how much replication was performed in both types of experiments. We selected experiments reporting differential expression under well-defined conditions and omitted those with more complex pooling or splitting protocols which complicate the accounting. Only single-color microarray studies were used, as being closest to RNA-seq in design (the dye-effect technical controls having no RNA-seq equivalent). From Array Express in the period 2008-2014 we identified 99 RNA-seq studies and 109 microarray studies meeting the criteria. A custom perl script was used to count, for each study, the number of biological replicates per unique variable combination (e.g. virus inoculated + 50 hours) for either a library type (i.e. single- or paired-end read), or array design. Then the average number of biological replicates per study, and by technology if more than one was used, was determined.

### MINSEQE compliance and consistency of data in three persistent data archives

To compare MINSEQE compliance across the three repositories we randomly chose 100 RNA-seq studies from each, focusing on gene expression studies and not *de novo* transcriptome assembly papers. To prevent duplication, only direct submissions were considered for SRA (i.e., SRA datasets that had been “brokered” via ArrayExpress or GEO were excluded from the SRA set and were counted only as ArrayExpress or GEO submissions). While each submission was assessed by one of the study authors, 25 studies from each set were assigned to two of the authors, to make sure that our assessment process itself was consistent and accurate.

During the assessment process the submissions were viewed through the interface of the source database. Their compliance with MINSEQE standards was assessed based on the presence/absence of the recommended annotation categories, as follows: (i) experiment aim and type of experiment (e.g. RNA-seq of coding RNA or RNA-seq of non-coding RNA), (ii) submitter’s contact details (iii) sample (target species) and experimental variable information (what varies between samples) (iv) information about biological and technical replicates, (v) experimental (e.g. treatment, library construction) and data processing (e.g. mapping) protocols, (vi) availability of raw sequencing reads and (vii) availability of processed data.

---

**Supplementary Table 1.** ICGC, ENCODE and Geuvadis guidelines comparison.

|  |  | ICGC guidelines for a gene expression experiment<sup>1</sup> | The ENCODE consortium, Standards, Guidelines and Best Practices for RNA-Seq V1.0<sup>2</sup> | Geuvadis consortium<sup>3</sup> |
| --- | --- | --- | --- | --- |
| Sample meta-data | Sample information | X | X | X |
|  | Identification of the type of RNA targeted |  | X | X |
|  | Protocols used to isolate RNAs |  | X | X |
|  | Methods used to quantify RNAs prior to sequencing |  | X | X |
| Experiment design | Replicate number |  | X |  |
|  | Sequencing depth |  | X | X |
| Sample preparation meta-data | Method of Preparation of cDNA | X<br>experimental_protocol | X | X |
|  | Quantitative standards (spike-ins) |  | X |  |
| cDNA and sequencing design | stranded or unstranded |  | X |  |
|  | pool and bar-coded RNA targets |  | X |  |
|  | sequencing platform used | X platform | X | X |
|  | depth of sequencing (e.g. number of reads obtained) |  | X | X |
|  | length of sequence reads |  | X | X |
|  | single or paired-end |  | X | X |
|  | base calling algorithm | X |  |  |
|  | sequencing strategy | X |  |  |
| Post-sequencing mapping, read statistics, quality scores: Mapping of sequence data | The program and version used, parameters employed, etc. | X<br>alignment_algorithm | X | X |
|  | The sequence version of the reference genome and, as appropriate, collections of transcript model sequences or splice junction | X<br>assembly_version<br>gene_build_version | X | X |
|  | Treatment of reads mapping equally well to more than one site in the reference genome |  | X |  |
|  | Mapping thresholds used |  | X |  |
|  | Specify if mapping was performed relative to the genome and/or transcriptome or a combination |  | X | X |
|  | Specify the order of steps in the mapping pipeline: simultaneous genome and transcriptome mapping, or stepwise mapping, etc. |  | X |  |
| Information concerning mapped results | Total number of uniquely mapped reads (i.e. occurs once in reference genome) |  | X |  |
|  | If paired-end reads are used, report the number of mapped read pairs and the number of mapped single reads |  | X |  |
|  | When appropriate, provide quantitation of various mapped elements including exons, splice sites, transcripts, etc. | X<br>normalization_algorithm | X |  |
|  | Reproducibility of replicates (e.g. IDR) |  | X |  |
|  | Analysis of spike-in standards |  | X |  |
| Raw sequence read data for each assay | FASTQ files | X |  | X |
| Final' processed (or summary) data for the set of assays in the study | BAM files | X |  | X |
|  | Normalized counts | X |  | X<br>analysis results |

<sup>1</sup>https://docs.icgc.org/submission/guide/intro/
<sup>2</sup>https://www.encodeproject.org/about/experiment-guidelines/
<sup>3</sup>http://www.internationalgenome.org/data-portal/data-collection/geuvadis

